# Detection and Spatiotemporal analysis of *in-vitro* 3D migratory Triple-Negative Breast cancer cells

**DOI:** 10.1101/2021.07.29.454312

**Authors:** Nikolaos M. Dimitriou, Salvador Flores-Torres, Joseph Matthew Kinsella, Georgios D. Mitsis

**Affiliations:** Department of Bioengineering, McGill University, Montreal, QC, H3A 0E9, Canada

**Keywords:** Cell segmentation, Point Pattern analysis, Confocal microscopy, 3D cell cultures

## Abstract

The invasion of cancer cells into the surrounding tissues is one of the hallmarks of cancer. However, a precise quantitative understanding of the spatiotemporal patterns of cancer cell migration and invasion still remains elusive. A promising approach to investigate these patterns are 3D cell cultures, which provide more realistic models of cancer growth compared to conventional 2D monolayers. Quantifying the spatial distribution of cells in these 3D cultures yields great promise for understanding the spatiotemporal progression of cancer. In the present study, we present an image processing and segmentation pipeline for the detection of 3D GFP-fluorescent Triple-Negative Breast Cancer cell nuclei, and we perform quantitative analysis of the formed spatial patterns and their temporal evolution. The performance of the proposed pipeline was evaluated using experimental 3D cell culture data, and was found to be comparable to manual segmentation, outperforming four alternative automated methods. The spatiotemporal statistical analysis of the detected distributions of nuclei revealed transient, non-random spatial distributions that consisted of clustered patterns across a wide range of neighbourhood distances, as well as dispersion for larger distances. Overall, the implementation of the proposed framework revealed the spatial organization of cellular nuclei with improved accuracy, providing insights into the 3 dimensional inter-cellular organization and its progression through time.

## 1 Introduction

An important aspect of cancer progression is the migration of cancer cells to the surrounding tissues. Both *in-vivo* and *in-vitro* studies on cancer cell migration have shown that cancers can exhibit several types of patterns including single cell migration, multicellular streaming and collective cell migration, as well as passive patterns, such as tissue folding, and expansive growth.^12^ Some of these patterns are found in invasive tumours such as breast cancer. Previous studies have shown that the tumour border of breast cancers is dominated by collective cell migration forming small acinar structures.^12^ Evidence of multicellular streaming also exist from orthotopic breast cancer in xenograft mouse models.^26^ Other clinical studies on the morphology of the surface of infiltrating ductal adenocarcinoma have shown that the fractal dimension of cancerous tissue is larger compared to normal breast tissue.^21^ Despite the fact that a significant amount of knowledge has been recently obtained for the qualitative characteristics of cancer invasion both *in-vivo* and *in-vitro*, there is still incomplete information regarding the quantitative characterization of cancer progression, and the investigation of the tumour organization.

To this end, 3D cell culture models have become a very promising experimental tool. The main reasons are the increased control of the experimental conditions, the flexibility of data collection compared to *in-vivo* experiments, and their more realistic representation of tumour progression compared to 2D cultures. Differences between 3D and 2D cultures have been observed in cancer growth and its related biochemical processes, such as the secretion of extracellular matrix (ECM) components, and intercellular interaction components^15^ while the histological and molecular features of *in-vitro* 3D spheroids exhibit more similarities with xenografts than the conventional 2D monolayers.^15^ Another advantage of 3D cell culture models is their flexibility with regards to incorporating more than one cell populations, such as stromal cells, as well as on changing the stiffness of the ECM. The heterotypic intercellular interactions between cancer cells and stromal cells, such as fibroblasts, results in altered cancer cell proliferation and migration, as well as the formation of more compact spheroids compared to equivalent 3D cell mono-culture systems. ^15^ Additionally, the collection of imaging data for *in-vitro* 3D cell cultures is generally easier and more accurate than *in-vivo* models. Intravital imaging is a common way of data collection for *in-vivo* models; however, this technique suffers from technical challenges such as passive drift of cells or tissues, low penetration depth, tissue heating, and limitations on imaging intervals.^12^ On the other hand, confocal microscopy used for *in-vitro* 3D cell cultures can produce higher resolution images, and the data collection intervals are more flexible. Although, 3D cell cultures cannot yet capture the full complexity of tumour growth in a living tissue, overall they have a lot to offer as they provide the opportunity to track even single cells.

Confocal microscopy of fluorescent cells, and cell segmentation algorithms are two important tools for the study of 3D *in-vitro* cancer growth. However, some common technical issues related to these two techniques may limit the tracking ability of cancer progression. Increased autofluorescence from out-of-focus cells, and variations in fluorescent signal intensity among the cells may pose challenges to cell segmentation algorithms resulting in over- or under-segmentation of cells.^11^ At the same time, the problem of image segmentation is ill-posed, and up to now there are no algorithms that can be considered as a gold standard. Some key algorithms developed for this purpose are intensity-^23^, boundary-^35^, region-based and region-growing.^24^ However, they present limitations on the distinction of individual cells in cases of uneven illumination, and in cases where the cells are in contact.

To account for this, algorithms that combine multiple methods have been proposed, thus improving the segmentation performance. ^19,32^ Watershed segmentation algorithms have become popular due to their capability to separate touching cells by utilizing information from their geodesic distance maps, even though they are prone to over-segmentation.^31^ Several variants of the watershed segmentation that address this issue have been proposed, with the marker controlled or seed based watershed transformation being the most popular.^28^ The separation of fused cells has also been approached with concavity-based techniques, which search for the optimal path between two concave points and separate the cells under the assumption that their fused shape contains concavities.^33^ More sophisticated energy minimization techniques^16^ have also been developed; however, their application to datasets containing a large amount of cells can be computationally prohibitive. Novel machine learning based methods exhibit improved performance, however their applicability may be limited to specific datasets, and their performance may be decreased in datasets with high cell shape and volume heterogeneity, as well as high cell density.^20^ Recent studies have mainly focused on the preprocessing of 3D image stacks, to improve the segmentation results of simpler segmentation algorithms, and are applicable to large datasets.^20^ Concluding, the advances made in both experimental and image processing methods provide us with the opportunity to further investigate the spatiotemporal organization and progression of cells in greater detail using 3D cell cultures.

Even though technological advances have provided us appropriate tools for a detailed and quantitative study of spatiotemporal cancer progression, our knowledge so far is rather limited to mostly qualitative aspects of this progression. The possibility of interpreting a cell as a point in the 3D space allows more quantitative, spatial statistical techniques to be employed.^14^ Spatial statistical techniques including the Complete Spatial Randomness (CSR) test^10^, the characterization of cell distributions using their Inter-Cellular and Nearest-Neighbour distances, and the analysis of cellular density profiles have already been applied for the investigation of tumour morphology and heterogeneity in histology images^7^, the whole-cell dynamic organization of lysosomes^1^, as well as the investigation of cell clustering and the correlation the genomic profile of tumours from tissue slices. ^34^

In this context, we present a processing, segmentation, and spatiotemporal analysis pipeline for the detection of *in-vitro* 3D cultured fluorescent cancer cells and investigation of their spatiotemporal progression. The proposed pipeline, presented in Fig. 1, utilizes a combination of preprocessing and segmentation algorithms to improve the detection performance of GFP-fluorescent nuclei of Triple Negative Breast Cancer (TNBC) cells. The performance of the proposed pipeline was evaluated against manual segmentation, and four alternative pipelines including established^8^, and novel machine learning algorithms.^20,30^ The segmented nuclei were subsequently used for the investigation of their spatiotemporal progression using point-pattern analysis, and density analysis methods. The novel combination of these methods enabled us to detect the position of the cells in the 3D space with higher accuracy, as well as to examine the organization and progression of cancer growth.

**Figure 1:**
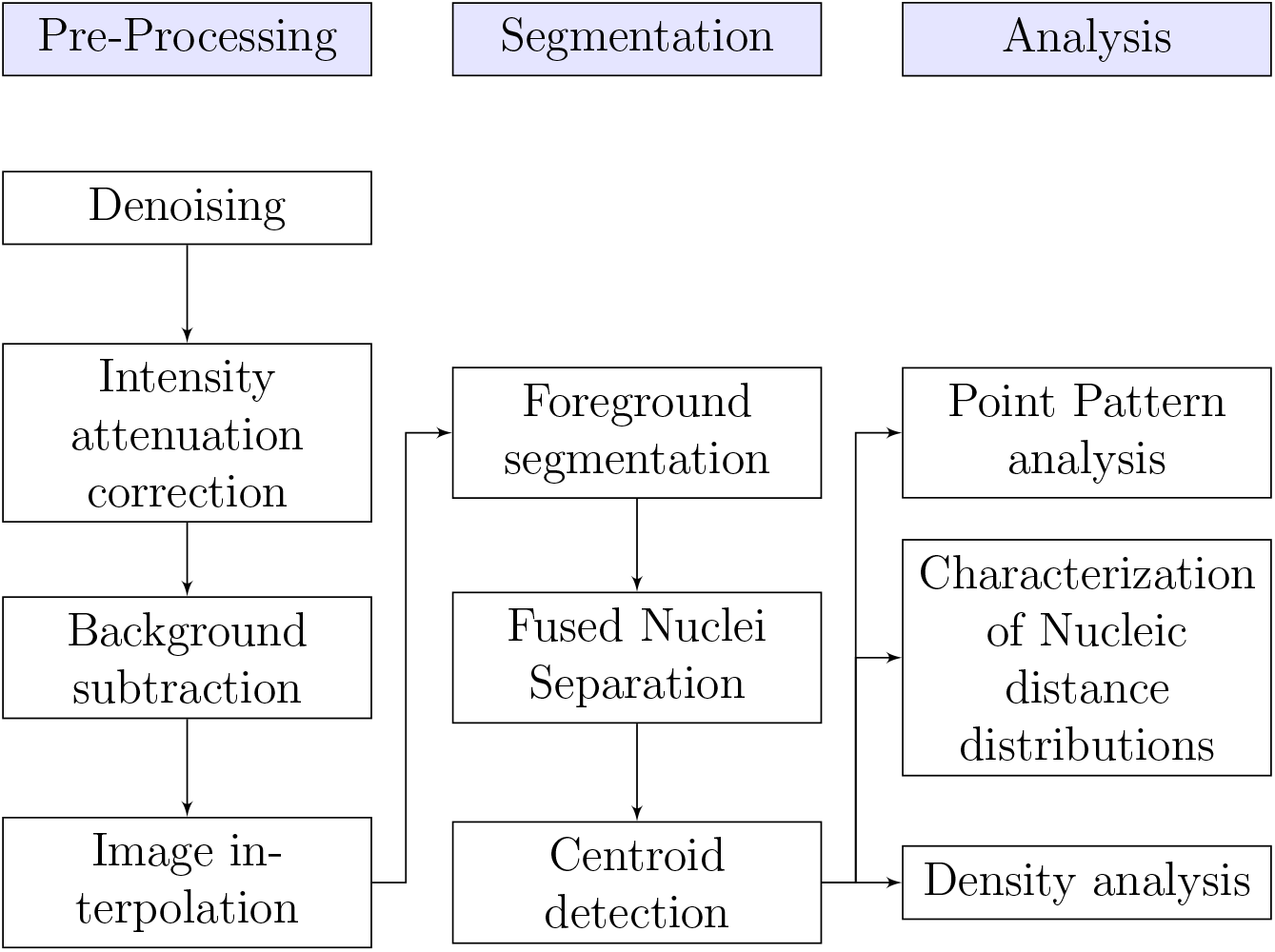
Proposed pipeline for the preprocessing of 3D image stacks, segmentation of fluorescent nuclei, and their spatiotemporal statistical analysis.

## 2 Materials and Methods

### 2.1 Experiments

#### 2.1.1 Cell preparation

TNBC cells from the MDA-MB-231 cell line with nuclear GFP (histone transfection), were thawed and cultured at 5% CO2, 37 °C in DMEM (Gibco) at pH 7.2 supplemented with 10% fetal bovine serum (Wisent Bioproducts), 100 U/mL penicillin, 100 μg/mL streptomycin, and 0.25 μg/mL, and amphotericin B (Sigma) in T-75 flasks (Corning). The cells were passaged before reaching 85% confluence. Three passages were performed before the 3D cultures; cells were rinsed twice with DPBS and trypsin-EDTA (0.25%-1X, Gibco) was used to harvest them.

#### 2.1.2 3D cell cultures

A cell-Matrigel (Corning) suspension was created using 0.25 mL of Matrigel (4 °C) and 5 × 10^4^ MDA-MB-231/GFP cells. Droplets of 5 μL cell-Matrigel mixture were manually deposited onto a high performance #1.5 glass bottom 6-well plate (Fisher Scientific). Data acquisition was performed using a confocal microscope (Nikon A1R HD25) coupled with a cell-culture chamber every 2-3 days for a total of 15 days. The dimensions of the 3D cultures were approximately 2500 × 2500 × 900 μm^3^. Cell localization was made possible by the GFP fluorophore that is present in cell nuclei (supplementary Fig. S.1). For this study 12 datasets were produced with samples from days 0, 2, 5, 7, 12, 14 each.

### 2.2 Image preprocessing

#### 2.2.1 Denoising

Poisson-noise is commonly found in low intensity fluorescent microscopy images.^17^ The selected denoising method was the *Poisson Unbiased Risk Estimation-Linear Expansion of Thresholds* (PURE-LET) technique implemented on ImageJ.^17,27^ This method is based on; the search of the closest possible noise-free signal by minimizing the unbiased estimate of the mean squared error (MSE) between the noise-free signal estimates and the noisy signal, the linearity of the estimates, and 3) the use of interscale predictors for the denoising process. The method utilizes a mixed Poisson-Gaussian noise model of the following form

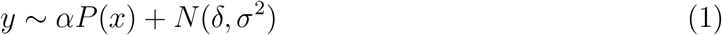

where *y* is the noisy input data, *x* the noise-free data, *P(x)* the Poisson-corrupted input data, *α* the detector gain, *δ* the detector offset, and *σ* the standard deviation of the additive white Gaussian noise. The estimation of the noise parameters (*α, δ, σ*) is fully automated.

#### 2.2.2 Intensity attenuation correction

Confocal microscopy image stacks are usually accompanied by decreasing intensity effects as the depth of the sample increases. The algorithm selected for the attenuation correction^4^ is implemented on ImageJ, and assumes a stationary background throughout an image stack and rectifies the average intensity and standard deviation by applying a linear transformation to each image slice according to

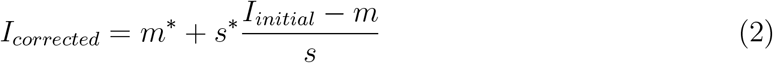

where *m\** and *s\** are the background average intensity and standard deviation of the reference slice respectively, and *m, s* are the background average intensity and standard deviation of the current slice. For each slice, the background is estimated by applying a morphological opening with a flat structuring element of radius equal to the radius of the smallest observed nucleus. The estimated background is then subtracted from the corrected image.

#### 2.2.3 Background subtraction

The background subtraction step during the correction of the attenuated intensity was found to be insufficient, because it subtracts only the local background around the nuclei. To fully eliminate background effects, we performed two additional steps. The first step was the background subtraction using the rolling ball algorithm.^29^ The rolling ball algorithm calculates a local background value for every pixel by averaging over a large ball surrounding the pixel. Using High-Low Look Up Tables (HiLo LUTs) we examined the background effects, and proceeded to manual thresholding of low intensity values, if these effects persisted. In our dataset this threshold was ~20 for 8-bit images, however its selection was performed separately for each image, due to varying intensity distributions across samples.

#### 2.2.4 Interpolation

Our sample consisted of images with resolution 999 × 999 pixels that corresponded to an area of around 2.5 × 2.5 mm^2^. A nucleus cross-section may have an approximate area of 130 μm^2^, which corresponds to a radius 6.43 μm, under the assumption that the cross-section is circular. This translates to a radius of 2.6 pixels. The small size of the nucleus may pose problems during the segmentation due to the fact the intensity gradients may be very steep and narrow. To account for this, we upscaled 10 times each slice of the image stack using cubic spline interpolation. The interpolation was performed in MATLAB. ^18^

### 2.3 Nuclei segmentation

#### 2.3.1 Foreground segmentation

The interpolated images were then used as the input for the Marker based Watershed segmentation algorithm. The foreground objects are marked and segmented, iteratively for each image-slice by performing the following steps (supplementary section 2.1): 1) Contrast-limited adaptive histogram equalization (CLAHE) on the initial interpolated grey-level image. 2) Erosion on the CLAHE image using a disk shaped mask of radius 10 pixels. 3) “Opening by reconstruction” on the CLAHE image using the markers of step 2 to identify high-intensity objects in the CLAHE image. 4) Dilation of the reconstructed image of step 3 using a disk shaped mask of radius 10 pixels. 5) “Closing by reconstruction” of the complement image of step 3 using the complement image of step 4 as marker. 6) Detection of regional maxima in the image of step 5. 7) Binarization of the image of step 5 using a locally adaptive threshold calculated by Bradley’s method.^6^ 8) Closing of the image of step 6 using a disk shaped mask of radius 5 pixels followed by morphological erosion, small object removal (with size less than 10 pixels), and filling of holes. 9) Gradient of the CLAHE image. 10) Imposed minima on the result step 9 using as mask the union of the complement image of step 7 and the image of step 8. 11) Watershed image of 10. The watershed map was then binarized^6^ and filtered to remove potentially small or very large artefacts that persisted. The algorithm was implemented in MATLAB.

#### 2.3.2 Fused Nuclei Separation & Centroid Detection

Fused nuclei usually exhibit overlapping intensity distributions that the Marker-Controlled Watershed transform cannot separate. Instead, their separation can be achieved by taking into account their morphological characteristics. In this step, we performed a classic distance based watershed segmentation on the Euclidean distance map of the nuclei detected by the afore mentioned Marker-Controlled Watershed algorithm. The centroids of the segmented nuclei were then detected by tracing the edges of the segmentation masks using a 26-connected neighbourhood tracing algorithm implemented in MATLAB.

### 2.4 Spatial Analysis

#### 2.4.1 Complete Spatial Randomness Test of nucleic distributions

The Complete Spatial Randomness (CSR) test examines whether the observed spatial point patterns can be described by a uniform random distribution.^9^ The CSR test was performed using Ripley’s *K*-function and the *spatstat* ^3^ package of R.^25^ The *K*-function^10^ is defined as the ratio between the number of the events, i.e. locations of points, *j* within a distance t from the event i, over the total number of events N, in the studied volume V (2.5×2.5×0.9 mm^3^)

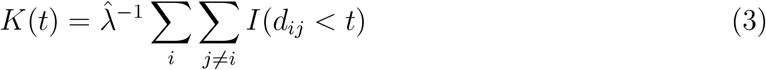

where 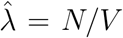 denotes the average density of events, *N*, in the studied volume *V*, *d*_*ij*_ is the distance between events *i* and *j, t* is the search radius and *I* a decision function

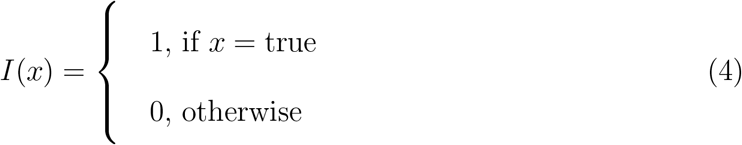

The *K*-function was calculated across all datasets and compared against complete spatial randomness that follows a Poisson process *K(t)* = 4πt^3^/3 in 3D.^10^ Isotropic edge correction was applied in the calculation of the *K*-function. To assess the uncertainty of the random variable *K* we produced a CSR envelope by generating 100 random distributions and calculating the *K*-function for each of them. The envelope was created by keeping minimum and maximum values of the resulted *K* values. A substantial upward separation of the observed *K*-function from the theoretical random *K*-function denotes clustered patterns, while a downward separation denotes dispersed patterns respectively. Both separation types suggest non-randomness in the distributions.

#### 2.4.2 Characterization of the Nucleic Distributions

##### Inter-Nucleic (IN) Distance Distributions

The IN Distance Distribution for a given sample was calculated by the pairwise Euclidean distances between all nuclei. Given two nuclei i and j with centroid positions **p_i_** = (*x*_*i*_, *y*_*i*_, *z*_*i*_) and **p_j_** = (*x*_*j*_, *y*_*j*_, *z*_*j*_) respectively, their pairwise Euclidean distance is given by 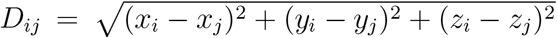 *i, j* = 1…*N*, *i* ≠ *j* where *N* the total number of nuclei. The similarity between two IN Distance Distributions of different t ime-points w as e stimated u sing t he c osine similarity measure (supplementary section 3).^13^

##### Nearest-Neighbour (NN) Distance Distributions

The NN Distance Distribution for a given sample was calculated using the distances between the nearest neighbours of the nuclei. The NN distance for a given nucleus *i* is given by the minimum IN Distance between the nucleus *i* and all the other nuclei of the sample, such as 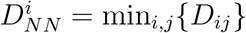, *j* ∈ [1, *N*], *j* ≠ *i*. Similarly, we used the cosine similarity measure to estimate the similarity between two NN distance distributions from different time-points. The IN, and NN Distances, as well as the similarity tests were computed in MATLAB.

#### 2.4.3 Density profiles

The CSR test and the characterization of the nucleic distance distributions can provide information on the structure of the spatial nucleic distributions. However, they do not provide sufficient in formation ab out th e lo cation of th ese di stributions in 3D sp ace. The final s tep o f the s patial analysis was the e xamination o f the regions w here c lustering takes place. To investigate the density of the nucleic distributions and their corresponding locations in 3D space, we estimated the density profiles of the centroids of the nuclei using the Kernel Density estimation via the Diffusion method.^5^

## 3 Results

### 3.1 Preprocessing, Segmentation, and Assessment of Performance

The detection of the fluorescent of TNBC nuclei cultured in 3D Matrigel ECM was performed using the proposed image preprocessing and segmentation pipeline. The preprocessing stage consists of denoising, intensity attenuation correction, background subtraction, and image interpolation, while the segmentation stage consists of foreground segmentation, splitting of fused nuclei, and detection of the centroids of the segmented nuclei. The effects of preprocessing and segmentation can be inspected in Fig. 2a–2e and Fig. 2f–2g. In Fig. 2h the segmentation result is then rescaled to the original size of the image.

**Figure 2:**
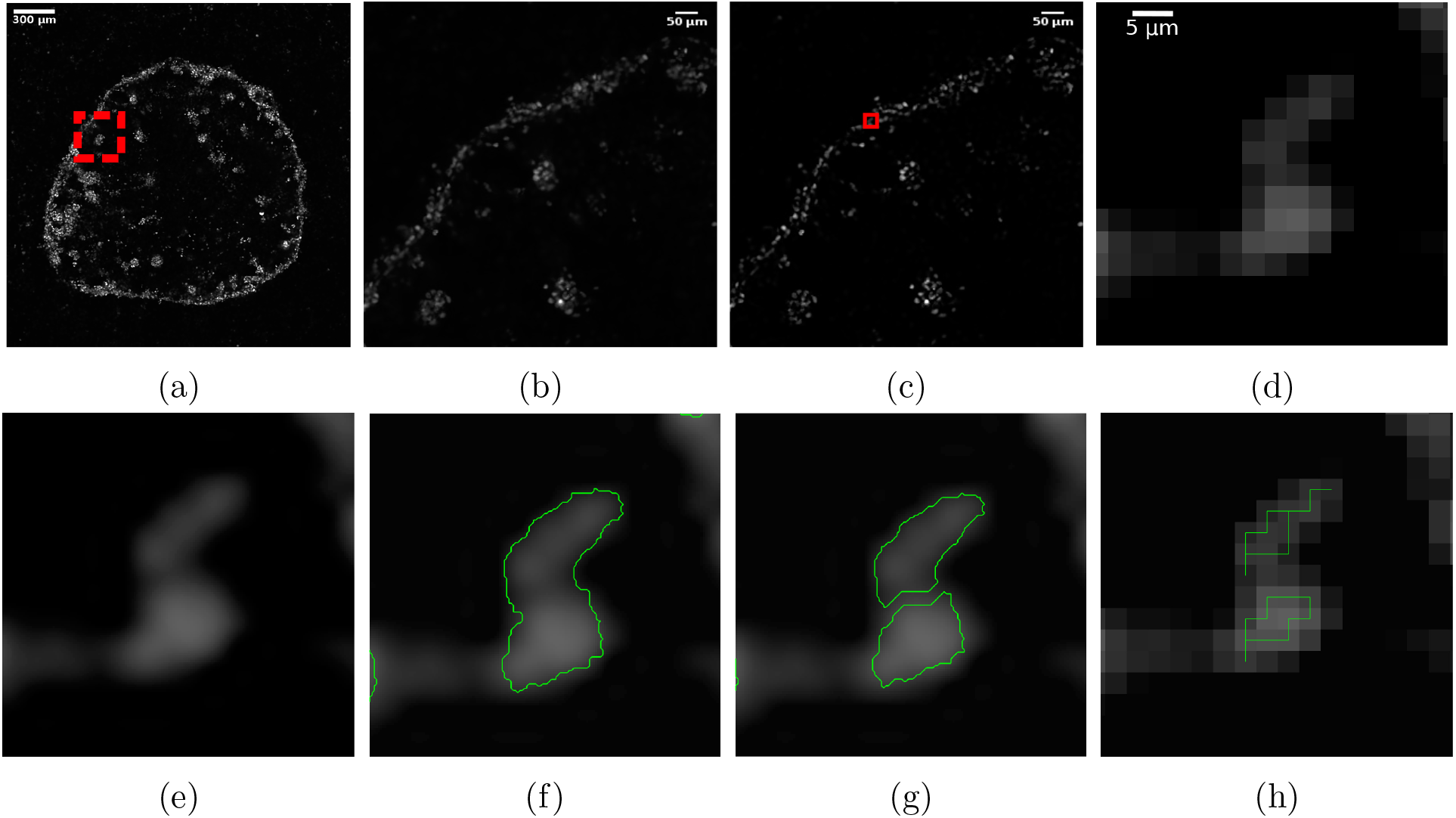
Segmentation process for fluorescent nuclei of 3D cell culture model. (2a) Raw image slice found at 80 μm height from the bottom of the well of a cell laden Matrigel dome in the 9th day of the experiment. (2b) Zoomed raw image of the red box region of (2a). (2c) Denoised and background subtraction result resulting from (2b). (2d) Zoomed image of the red box in (2c). (2e) Interpolation result resulting from (2d). (2f) Marker Controlled Watershed segmentation. (2g) Nuclei splitting with Distance Based Watershed segmentation. (2h) Rescaling back to original image size.

The detected centroids across the 7 time-points, as well as the average and standard deviation of the total number of nuclei across all datasets are depicted in Fig. 3, and supplementary Fig. S.2. The results show a biased movement of the cells towards the bottom of the plate. Furthermore, the cells exhibit a sigmoidal proliferative characteristic with numbers ranging from 1000 to 15000 nuclei.

**Figure 3:**
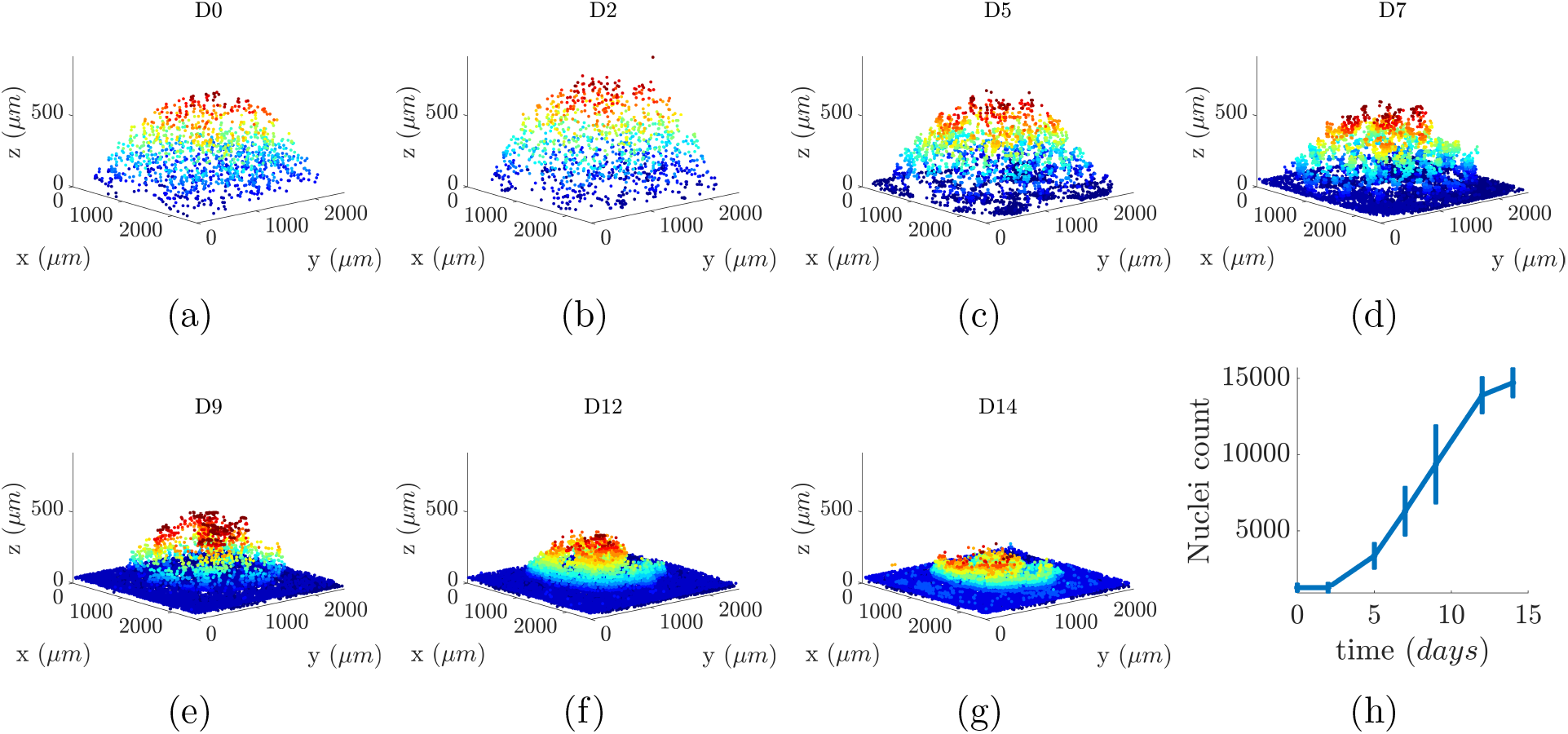
(3a)-(3g) Result from the application of the processing and segmentation pipeline in a representative dataset. Title notation D# refers to the time-point of the image acquisition in days. (3h) Mean and standard deviation for the nuclei count across 12 datasets.

The performance of the proposed preprocessing and segmentation pipeline (FluoDeSeg) was assessed using manual segmentation, by drawing the approximate borders of the nuclei, and compared to the performance of four alternative methods of Nasser et al. ^20^, a CellProfiler pipeline^8^, and two pretrained models (cyto and nuclei) of the Cellpose deep-learning segmentation algorithm.^30^ The segmentation performance was calculated using the accuracy, recall, precision, F1 score, Jaccard index (supplementary section 2.3). Additionally, the number of the segmented nuclei from all methods were also compared. Our method exhibited comparable accuracy compared to the manual annotation, and the highest accuracy, precision, F1 score, and Jaccard index among the three methods, as depicted in the summarized statistics of Table 1 and supplementary Fig. S3a.

**Table 1:**
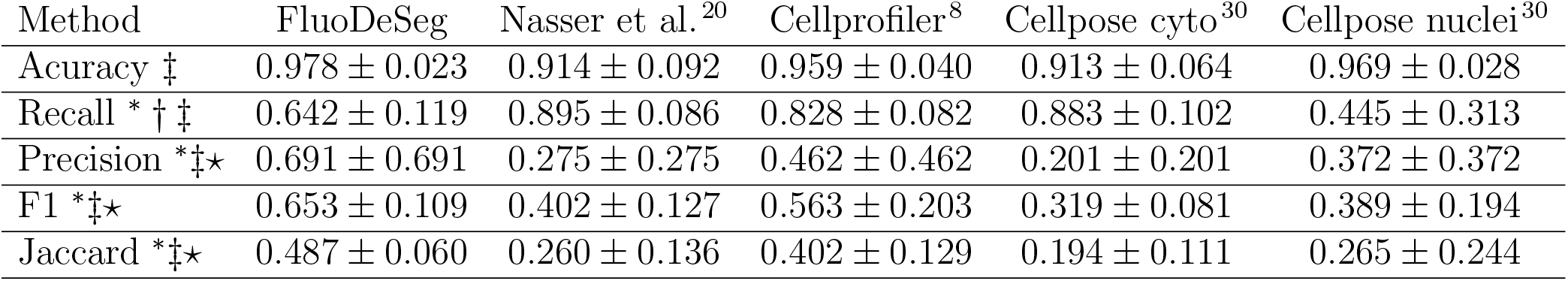
Segmentation performance of the proposed pipeline (FluoDeSeg), the method developed by Nasser et al.^20^, the CellProfiler pipeline^8^, and the two Cellpose models (cyto and nuclei)^30^ as compared to manual segmentation. Results are reported as Mean Standard Deviation. The symbols *, †, ‡, and ⋆ denote p-value < 0.05 for the Kruskal-Wallis test^22^ between FluoDeSeg and Nasser et al., between FluoDeSeg and CellProfiler, FluoDeSeg and Cellpose cyto, and FluoDeSeg and Cellpose nuclei, respectively.

### 3.2 Spatial Analysis

For the investigation of the spatial organization of the cells, we performed the CSR test, using Ripley’s *K-*function^10^, to examine whether the fluorescent nuclei, are randomly distributed in space. The results depicted in Fig. 5a indicate substantial differences from a uniform random distribution. Specifically, we observe clustering for a wide range of neighbourhood radii, as well as an increasing dispersion for longer distances across all samples, with respect to time.

The quantitative characterization of the spatial distribution of the cells was performed using the IN, and the NN Euclidean distance distributions. The IN distance distributions quantify the positioning of the cells relative to one another, while the NN distributions measure the distances between each cell and their nearest neighbouring cell. The resulting IN distance distributions, depicted in Fig. 5b, show that they remain relatively stable across all samples and time, with a characteristic peak distance at ~1 mm. The cosine similarity test yielded an average similarity value equal to 0.9946 ± 0.0074, suggesting high similarity between two IN distance distributions across different time-points. Their similarity remained high across all their time-point intervals as shown in supplementary Fig. S.3c. On the other hand, the NN distance distributions, presented in Fig. 5c, formed initially wide distributions that gradually tended to become narrower around lower neighbourhood radii values with respect to time, across all samples, with a characteristic peak at ~15 μm. The average cosine similarity between two NN distance distributions from different time-points was found to be equal to 0.8447 ± 0.1686. The similarity between two NN distance distributions was found to decrease as a function of the time separation between them, as shown in supplementary Fig. S.3d.

For the examination of the regions where clustering takes place, we estimated the density profiles of the centroids, using the Kernel Density Estimation via the Diffusion method^5^. The resulting density profiles, depicted in Fig. 6a–6g, suggest that cells were organized into clusters and these clusters tended to change positions in space with respect to time.

## 4 Discussion

In this study, a novel preprocessing and segmentation pipeline allowed us to examine the quantitative aspects of the spatiotemporal progression of cancer cells grown in cultured 3D Matrigel ECM. Based on this experimental setting, a more accurate detection of the fluorescent nuclei was achieved, compared to alternative segmentation methods. The spatial analysis revealed a dynamic behaviour of the detected nuclei across time, forming both clustered and dispersion patterns.

The pipeline was able to detect more accurately the fluorescent nuclei compared to the four examined alternative methods, and achieved comparable accuracy to the manual annotation. Specifically, our method achieved the highest accuracy, precision, F1 score, and Jaccard index score among the four methods (Table 1, supplementary Fig. S.3a). The lower recall score was due to an increased amount of pixels classified as False Negative. This result may be due to the background subtraction, which narrows the intensity distribution around the nuclei, even though the information about the location of the nuclei may not be lost. The comparison of the nuclei count of the five methods against the manual annotation showed that our method exhibited the best performance among all examined methods, with a maximum over-segmentation of around 5%, and a maximum under-segmentation of 14% as compared the nuclei count of the manual annotation (Fig. 4). The pipeline was applied to image stacks with planar resolution of 999 × 999 pixels that correspond to 2.5 × 2.5 mm^2^, and a nucleus radius of ~2.6 pixels. In most cases, the quality of the image stacks was sufficiently good to visually detect fused nuclei from their cluster. However, a possible decrease of the image resolution would most probably affect the segmentation performance. As a result, it is expected that the configuration of the microscope can have a significant effect on the achieved segmentation performance.

**Figure 4:**
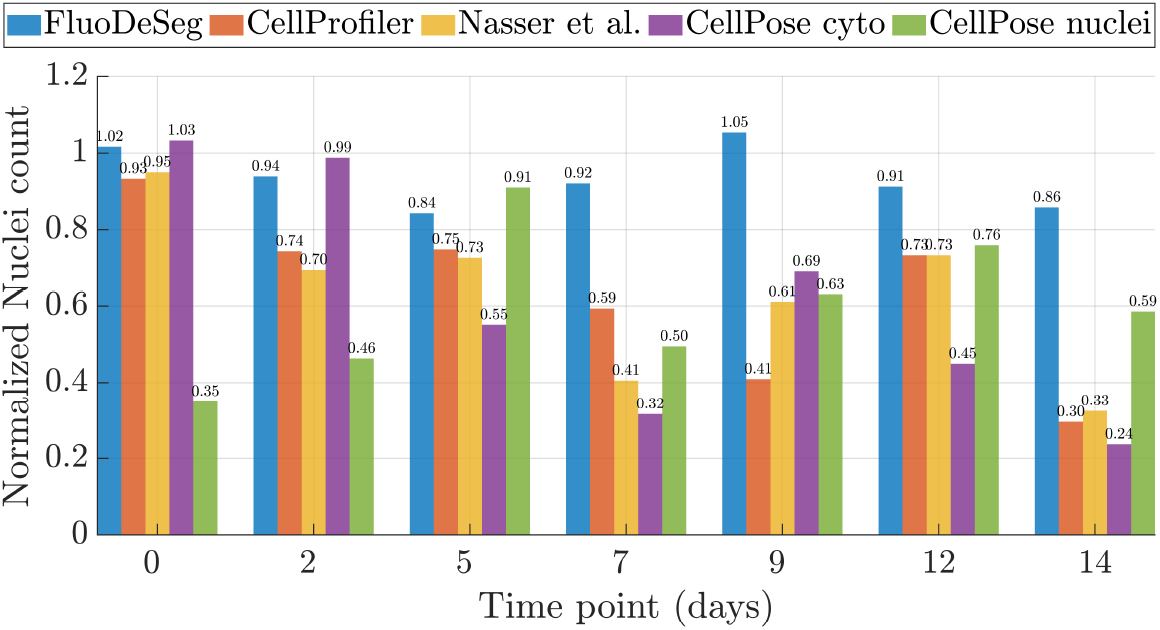
Normalized nuclei counts obtained by the three examined methods with respect to the nuclei count achieved by the manual annotation. Values greater than 1, and less than 1 denote over-segmentation, and under-segmentation, respectively. The results overall suggest that the proposed pipeline exhibits improved performance with respect to all the examined measures, with the exception of the recall score.

The CSR test revealed that the nuclei maintained clustered patterns for a wide range of neighbourhood radii across time and that they exhibited more pronounced dispersion patterns with respect to time (Fig. 5a). The biased movement of the cells towards the bottom may have contributed to the increase of clustering patterns in smaller neighbourhood radii. Even though Ripley’s *K-*function provides a measure of the formed patterns, a limitation of this measure is its insensitivity to different point patterns. Specifically, two different point patterns may result in the same *K-*function^2^. Thus, further steps had to be performed to extract more information about the formed patterns and their behaviour.

**Figure 5:**
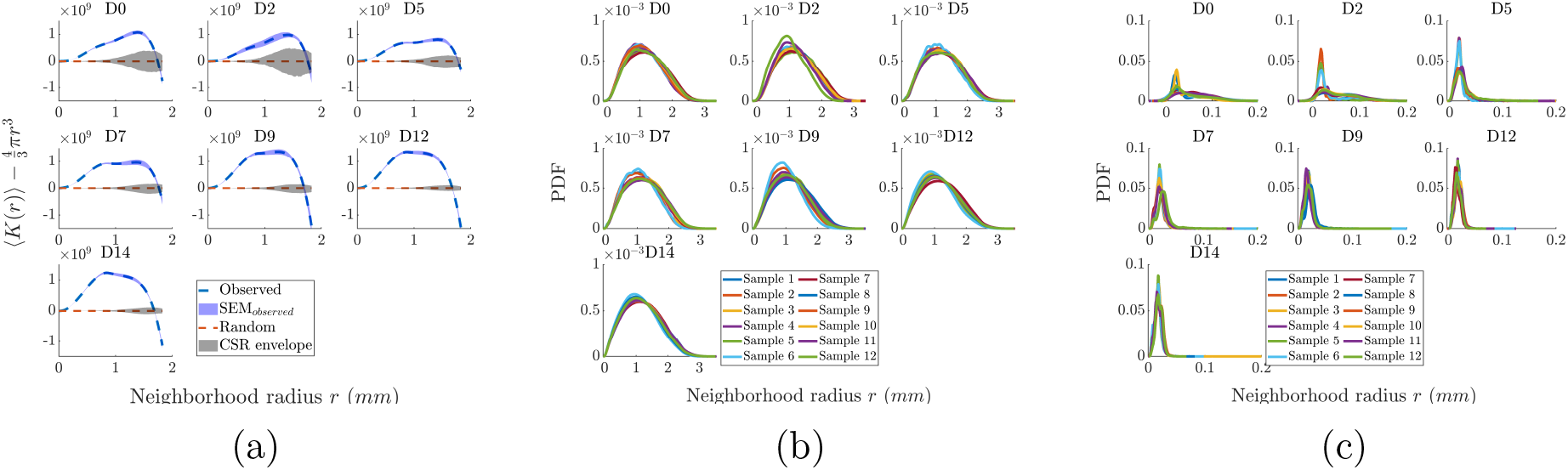
(5a) Average values of *K*-function across all samples and the corresponding standard error of mean (SEM). Upward separation of the observed *K*-function from the theoretical random *K*-function denotes clustered patterns, while downward separation denotes dispersed patterns. (5b) Inter-Nucleic Distance Distributions across all samples. (5c) Nearest-Neighbour Distance Distributions across all samples. The title (D#) denotes the time-point in days.

To investigate the regions of clustering and dispersion, we estimated the density profiles of the centroids of the nuclei. The results revealed organization of cells into smaller clusters and a dynamic behaviour of them in time with lower clustering regions appearing not only at the edges but also close to the center of the space (Fig. 6a–6g). These results, in combination with the results of the CSR test suggest that dispersed patterns did not only appear at the borders of the space, but also within its inner regions of it. This dynamic behaviour can be interpreted as a possible consequence of the need for balance between adhesiveness and access to nutrients, oxygen. While, it is crucial for cells to stay attached to each other, cell crowding may compromise their survival in the inner core of a cluster, due to limited diffusion of nutrients, oxygen, and accumulation of toxic metabolic waste products. We are currently investigating this behaviour by incorporating mathematical models.

**Figure 6:**
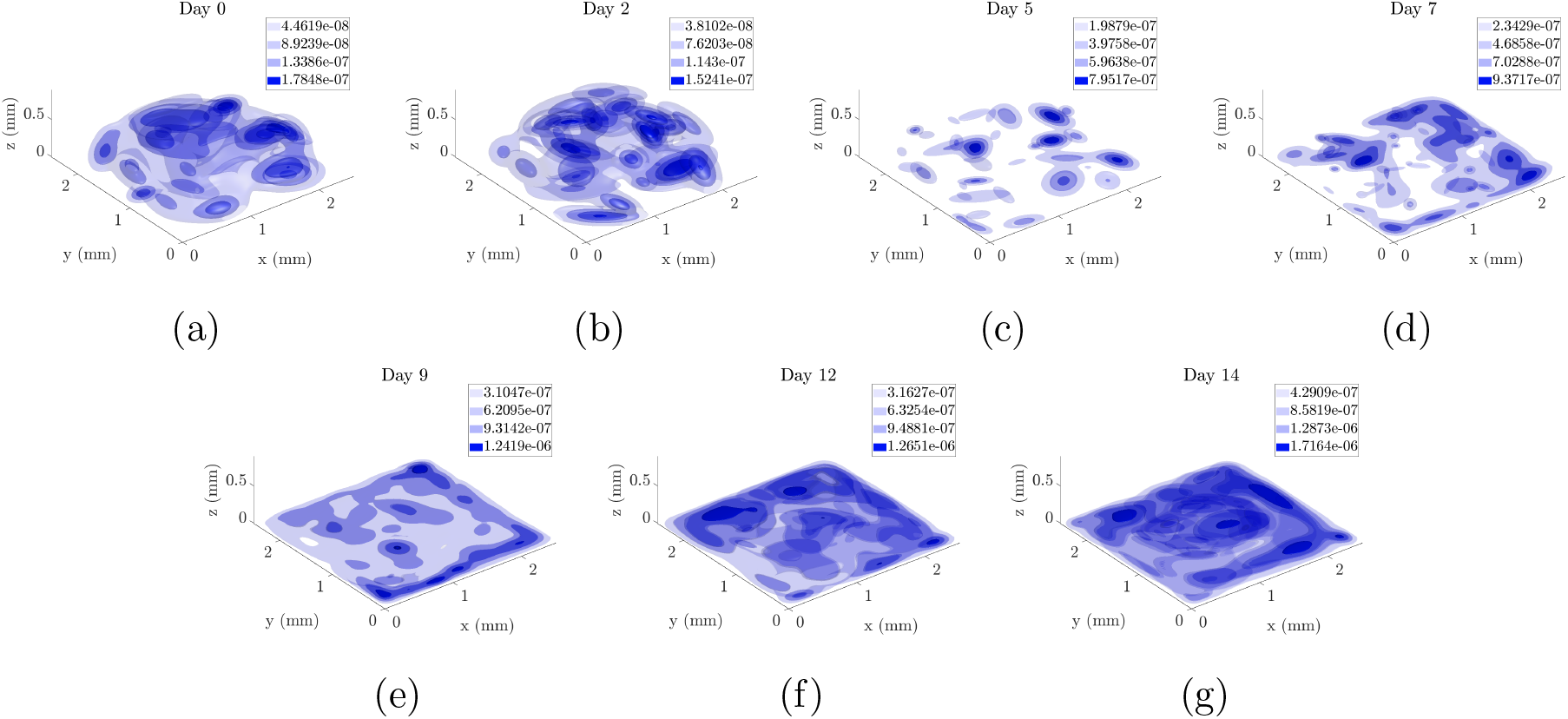
(6a)-(6g) Density profiles of a representative sample across time. The values in the legends indicate the density values of the contours painted with the same colour.

The IN distance distributions remained stable across all samples (Fig. 5b), maintaining high similarity for different time-points (supplementary Fig. S.3c), and the NN distances were initially widely distributed, and tended to become more narrow around smaller neighbourhood radii across time (Fig. 5c), with a decreasing similarity as the distance between time-points increases (supplementary Fig. S.3d). Although we would expect that the increasing reduction of the NN distances of these distributions would destabilize the IN distance distributions, this was not found to be the case. The maintenance of the stability of these distributions can be interpreted as a result of the organization of cells into clusters, the fact that cells tended to concentrate towards the bottom of the space with respect to time, as well as their synchronized division.

Concluding, the improved performance of the proposed pipeline compared to alternative methods allowed further quantitative investigation of the spatiotemporal progression of these cells. The employed spatial statistical methods allowed the extraction of information for the behaviour of cells across space and time, and the total tumour organization. Future directions include the application of the proposed framework to the investigation of *in-vitro* models with increased complexity, including the incorporation of stromal cell populations in 3D cell cultures, as well as the validation of more sophisticated spatiotemporal mathematical models using 3D cell culture data.

## Acknowledgment

N. M. D. thanks Stavros Niarchos Foundation (F237055R00), Werner Graupe (F202955R00) and McGill University (90025) for the scholarships. S. F. T. thanks McGill University for the McGill Engineering Doctoral Award (90025) and the FRQNT (291010) for the scholarships. This work was supported by Cyprus Research and Innovation Foundation (Project: INTERNATIONAL/OTHER/0118/0018).

## List of Figures

1. Proposed pipeline for the preprocessing of 3D image stacks, segmentation of fluorescent nuclei, and their spatiotemporal statistical analysis. … … … . 21
2. Segmentation process for fluorescent nuclei of 3D cell culture model. (2a) Raw image slice found at 80 μm height from the bottom of the well of a cell laden Matrigel dome in the 9th day of the experiment. (2b) Zoomed raw image of the red box region of (2a). (2c) Denoised and background subtraction result resulting from (2b). (2d) Zoomed image of the red box in (2c). (2e) Interpolation result resulting from (2d). (2f) Marker Controlled Watershed segmentation. (2g) Nuclei splitting with Distance Based Watershed segmentation. (2h) Rescaling back to original image size. … … … … . . 22
3. (3a)-(3g) Result from the application of the processing and segmentation pipeline in a representative dataset. Title notation D# refers to the time-point of the image acquisition in days. (3h) Mean and standard deviation for the nuclei count across 12 datasets. … … … … … … … . 23
4. Normalized nuclei counts obtained by the three examined methods with respect to the nuclei count achieved by the manual annotation. Values greater than 1, and less than 1 denote over-segmentation, and under-segmentation, respectively. The results overall suggest that the proposed pipeline exhibits improved performance with respect to all the examined measures, with the exception of the recall score. … … … … … … … … … 24
5. (5a) Average values of *K*-function across all samples and the corresponding standard error of mean (SEM). Upward separation of the observed *K*-function from the theoretical random *K*-function denotes clustered patterns, while downward separation denotes dispersed patterns. (5b) Inter-Nucleic Distance Distributions across all samples. (5c) Nearest-Neighbour Distance Distributions across all samples. The title (D#) denotes the time-point in days. … … … … … … … … … … … … … 25
6. (6a)-(6g) Density profiles of a representative sample across time. The values in the legends indicate the density values of the contours painted with the same colour. … … … … … … … … … … … . . 26

## List of Tables

1. Segmentation performance of the proposed pipeline (FluoDeSeg), the method developed by Nasser et al.^20^, the CellProfiler pipeline^8^, and the two Cellpose models (cyto and nuclei)^30^ as compared to manual segmentation. Results are reported as Mean ± Standard Deviation. The symbols *, †, ‡, and ⋆ denote p-value < 0.05 for the Kruskal-Wallis test^22^ between FluoDeSeg and Nasser et al., between FluoDeSeg and CellProfiler, FluoDeSeg and Cellpose cyto, and FluoDeSeg and Cellpose nuclei, respectively. … … … … … … 28

## Footnotes

1 https://figshare.com/projects/3D-GROWTH-MDA-MB-231-SERIES-12/118989

2 https://github.com/NMDimitriou/3D-Preprocessing-Nuclei-Segmentation.git

3 https://github.com/NMDimitriou/3D-spatial-analysis-cell-nuclei.git

