## Supplementary Material for "Detection and Spatiotemporal analysis of *in-vitro* 3D migratory Triple-Negative Breast cancer cells"

#### 1 Experiment

In figure S.1, image slices were obtained at a distance of $100-120$ $\mu m$ from the bottom of the plate, with an imaging window of $2500\times 2500$ $\mu m^2$ and resolution $999\times 999$ pixels (6 of the 12 datasets are presented).

#### 2 Nuclei segmentation

##### 2.1 Foreground segmentation

Often, in watershed segmentation a gradient of a grey-level image is considered so that the boundaries of the objects are located at high gradient points [2]. One can imagine this gradient image as a topographic landscape of ridges and valleys. The signal of a cell or an aggregate of cells changes locally the gradient image and creates "catchment basins". A catchment basin comprises of all points whose path of steepest descent terminates at a local minimum. The water floods the catchment basins and watersheds separate the basins from each other. In the case of noisy data, multiple local minima can be imposed, resulting to over-segmentation. The marker-controlled watershed algorithm marks the foreground objects to suppress unwanted local minima [1].

The opening by reconstruction is a process in which the peaks of the eroded marker image identify the location of objects to emphasize on the mask image. The marker image dilates inside the mask image until the contour of the marker fits under the mask image.

Similarly, the closing by reconstruction is the process in which the peaks of the complement of a dilated image function as markers to identify the location of objects to emphasize on the complement of the mask image used for the opening by reconstruction. For this process, the marker image dilates inside the the complement-mask image until the contour of the marker fits under the complement-mask image [2].

#### 2.2 Centroids of the segmented nuclei

The centroids of the segmented nuclei for 6 out of 12 datasets and their corresponding growth curves are depicted in Fig. S.2.

#### 2.3 Assessment of Performance

The testing dataset consisted of 7 randomly selected parts of the image samples that correspond to different time points with resolution ranging between $100 \times 100$ and $250 \times 250$ pixels, with 3-4 image slices each, and nuclei count from 60 to 229.

The segmentation performance was calculated using the accuracy, recall, precision, F1 score, Jaccard index defined as follows

$$\text{accuracy} = \frac{\text{TP} + \text{TN}}{\text{TP} + \text{TN} + \text{FP} + \text{FN}} \tag{1}$$

$$\text{recall} = \frac{\text{TP}}{\text{TP} + \text{FN}} \tag{2}$$

$$\text{precision} = \frac{\text{TP}}{\text{TP} + \text{FP}} \tag{3}$$

$$\text{F1}_{\text{score}} = 2 \times \frac{\text{recall} \times \text{precision}}{\text{recall} + \text{precision}} \tag{4}$$

$$\text{Jaccard}_{\text{index}} = \frac{|\text{MASK}_{\text{manual}} \cap \text{MASK}_{\text{auto}}|}{|\text{MASK}_{\text{manual}} \cup \text{MASK}_{\text{auto}}|} \tag{5}$$

where TP, TN, FP, FN are the true positive, true negative, false positive, and false negative sum of pixels between the corresponding segmentation method and manual segmentation, respectively.

### 3 Spatial analysis

The cosine similarity measure emerges from the Euclidean dot product, in which for two given vectors $\vec{x}$, $\vec{y}$, their similarity is defined as $\text{sim}(\vec{x}, \vec{y}) := \cos(\vec{x}, \vec{y}) = \frac{\vec{x} \cdot \vec{y}}{||\vec{x}||_2 ||\vec{y}||_2}$ where $|| \cdot ||_2$ the Euclidean norm. The similarity measure can take values from -1 to 1 indicating exactly opposite, and exactly same vectors, respectively. If similarity is equal to zero then the vectors are orthogonal. Values of similarity between 0 and 1 denote low, intermediate and high similarity.

##### 3.1 Density profiles of the centroids

The produced density profiles of the centroids are presented in Fig. S.4.

#### 4 CellProfiler pipeline

The CellProfiler pipeline used for the comparison with the proposed pipeline contains the following modules; Correct Illumination (Gaussian filter, smoothing filter size: 10 px), Identify primary objects (Typical diameter of nuclei: 5-15 px, default settings that include global threshold using the minimum cross entropy method, and distinction of clumped objects using information from the intensity), Shrink objects by 1 px, Split objects (default settings), Convert objects to image (saved binary mask).

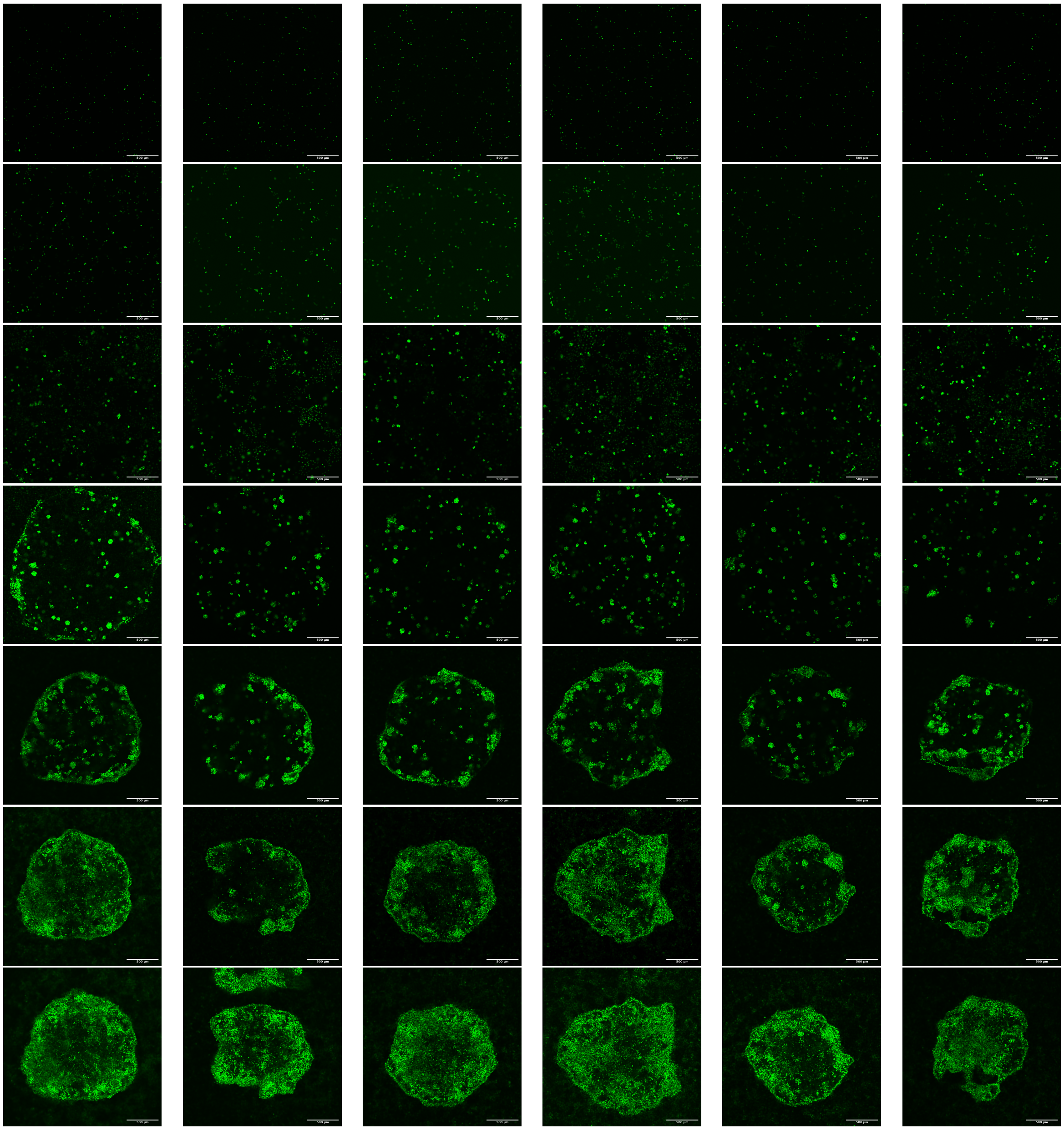

Figure S.1: Slices at $Z \approx 100\ \mu m$ from the bottom of the plate for 6 datasets across different time points. Each column represents a different dataset, and each row a different time point, namely day 0, 2, 5, 7, 9, 12, 14, from top to bottom. The images represent slices obtained at $Z \approx 100\ \mu m$ from the bottom of the plate. Scale bar: 500 $\mu m$.

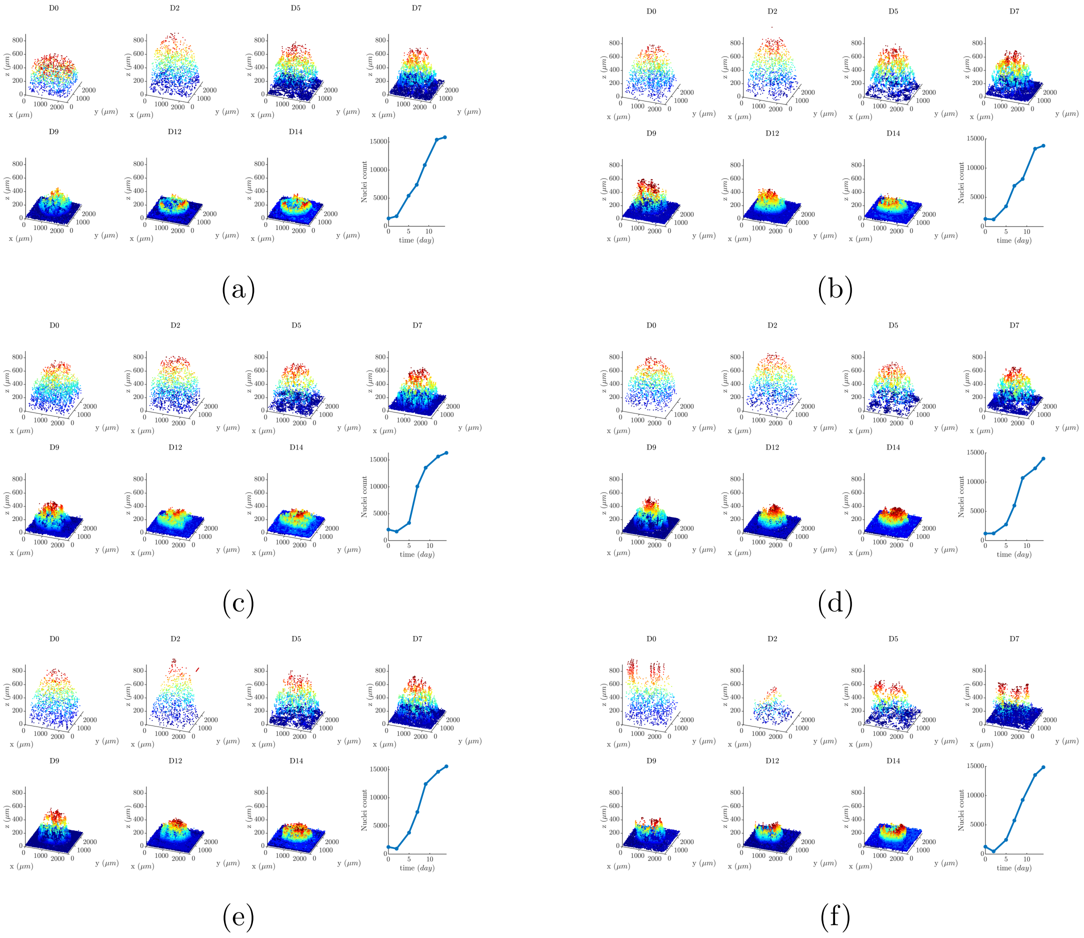


Figure S.2: Distribution of centroids of the segmented nuclei, and nuclei count with respect to time (6 out of the 12 datasets are presented).

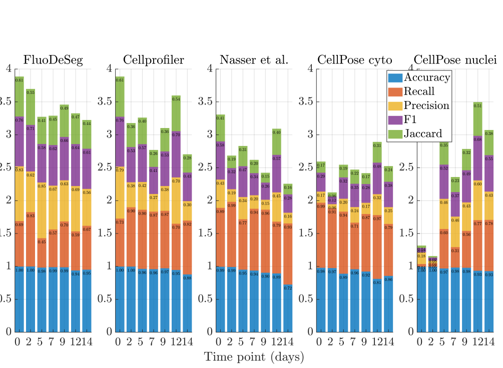


(a)

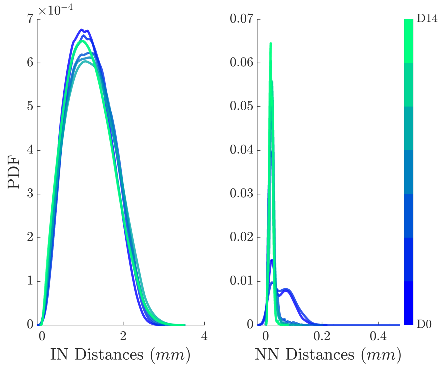


(b)

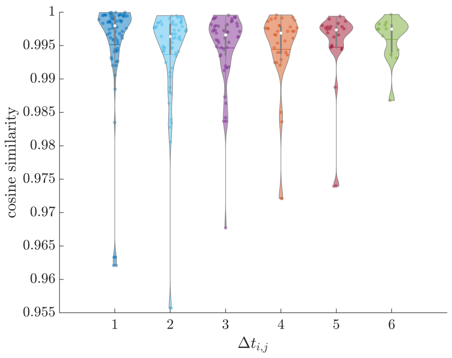


(c)

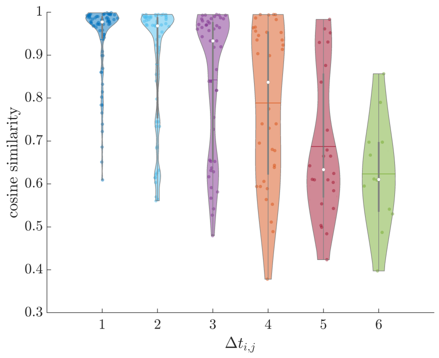


(d)

Figure S.3: (S.3a) Performance of the segmentation methods for samples at different time points. The summarized results of the performance metrics are presented in Tab. 1 of the manuscript. (S.3b) Inter-Nucleic and Nearest-Neighbour distance distributions of a representative sample colour-coded based on the time-point. (S.3c) Violin plots of cosine similarity with respect to different time-point intervals for Inter-Nucleic distance distributions. The Inter-Nucleic distance distributions remained highly similar across all time-points. (S.3d) Violin plots of cosine similarity with respect to different time-point intervals for Nearest-Neighbour distance distributions. The similarity between two Nearest-Neighbour distance distributions decreased for longer time separation. Note that for the $x$-axis, the values ranging from 1 to 6 indicate the result of the difference between the indices, $i$, $j$, of two time-points $t_i$, $t_j$. For example the value 6 is the result of the difference between $i = 7$ and $j = 1$ that correspond to $t_7 = 14^{\text{th}}$ and $t_1 = 0^{\text{th}}$ days, respectively.

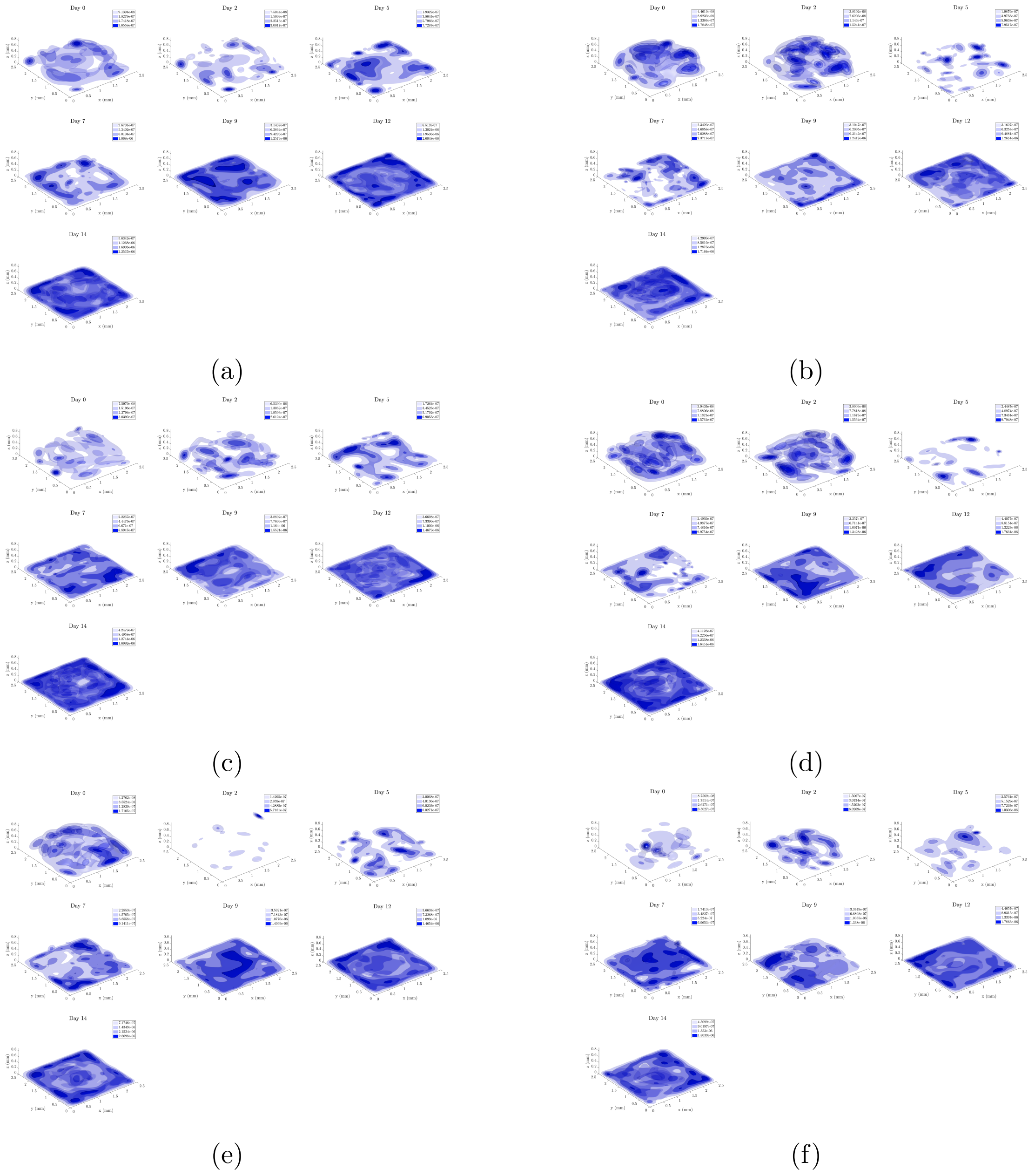


Figure S.4: Density profiles of the centroids across all time-points (6 out of the 12 datasets are depicted).
